# Mitogen-activated protein kinases associated sites of tobacco *REPRESSION OF SHOOT GROWTH* regulates its localization in plant cells

**DOI:** 10.1101/2021.11.16.468869

**Authors:** Luyao Wang, Ying Gui, Bingye Yang, Fangjie Si, Jianhua Guo, Chunhao Jiang

**Affiliations:** Department of Plant Pathology, College of Plant Protection, Nanjing Agricultural University, 210095 Nanjing, China; Key Laboratory of Integrated Management of Crop Disease and Pests, Ministry of Education / Key Laboratory of Integrated Pest Management on Crops in East China, Ministry of Agriculture/Key Laboratory of Plant Immunity, Nanjing Agricultural University, 210095, Nanjing, China; Engineering Center of Bioresource Pesticide in Jiangsu Province, 210095, Nanjing, China; Shenzhen Branch, Guangdong Laboratory of Lingnan Modern Agriculture, Genome Analysis Laboratory of the Ministry of Agriculture and Rural Affairs, Agricultural Genomics Institute at Shenzhen, Chinese Academy of Agricultural Sciences, Shenzhen, Guangdong 518120, China

## Abstract

Plant defense and growth rely on multiple transcriptional factors (TFs). REPRESSION OF SHOOT GROWTH (RSG) is known as one of the important TFs in tobacco (*Nicotiana tabacum*) with a basic leucine zipper domain. RSG was involved in plant gibberellin feedback regulation by inducing the expression of key genes. The tobacco calcium-dependent protein kinase, CDPK1 was reported to interact with RSG and manipulate its intracellular localization by phosphorylating Ser-114 of RSG. Here, we identified tobacco mitogen-activated protein kinase 3 (NtMPK3) as a RSG interacted protein kinase. Mutation of predicted MAPK-associated phosphorylation site of RSG (Thr-30, Ser-74 and Thr-135) significantly altered the intracellular localization of NtMPK3-RSG interaction complex. Nuclear transport of RSG and its amino acids mutants (T30A and S74A) were observed after treated with plant defense elicitor peptide flg22 in 5 min, while the two mutated RSG swiftly relocalized in tobacco cytoplasm in 30 min. Moreover, triple points mutation of RSG (T30A/S74A/T135A) mimics constant unphosphorylated status, and predominantly localized in tobacco cytoplasm. RSG (T30A/S74A/T135A) showed no relocalization effect under the treatments of either flg22, *B. cereus* AR156 or GA_3_, and was impaired in its role as TFs. Our results suggest that MAPK associated phosphorylation sites of RSG regulate its localization in tobacco and constant unphosphorylation of RSG in Thr-30, Ser-74 and Thr-135 keeps RSG predominantly localized in cytoplasm.

## Introduction

Developmental plasticity is a unique features of plant, and multiple complex programs were evolved to adapt to different environmental situations. Plant hormones are known to contribute to innate transcriptional processes in specialized manners and to transfer exterior environmental provokes to plant nuclei. Gibberllins (GAs) are tetracyclic diterpenoid phytohormones, which regulate plenty aspects of plant growth and development stages, such as seed germination, stem elongation, leaf expansion, flowering and fruiting (Matsuoka, 2003). In previous studies, REPRESSION OF SHOOT GROWTH (RSG) has been characterized as a transcriptional activator in tobacco, which was identified as basic leucine zipper (bZIP) family protein and regulate endogenous amounts of GAs through controlling the expression of GA biosynthetic enzyme (Fukazawa et al., 2010). RSG was also reported to interact with 14-3-3, which forms highly conserved family of homodimeric and heterodimeric protein complex (Ishida et al., 2004). It was suggested that 14-3-3 signaling proteins have been suggested to suppress RSG by restricting it in cytoplasm of tobacco cells and phosphorylation on Ser-114 of RSG was proven to be important for its binding with 14-3-3 (Ishida et al., 2004). Mutation of Ser-114 to Ala in RSG blocks its interaction with 14-3-3, and this mutated version of RSG (S114A) showed predominant localization in the nucleus after GA application (Ishida et al., 2004). A tobacco calcium-dependent protein kinase (CDPK1) was reported to decode calcium signal produced by GAs in tobacco and regulates intracellular localization of RSG. It is also suggested that CDPK1 was identified as a RSG kinase, which facilitates interaction between RSG and 14-3-3 (Ishida et al., 2008).

MAPKs are involved in highly conserved signaling pathways in eukaryotes, and studies on plant MAPKs have attracted great attentions as MAPKs were key signal mediators in plant during response to different environmental stresses (Andreasson and Ellis, 2010, Cheong and Kim, 2010, Sinha et al., 2011, Meng and Zhang, 2013). In previous reports, triggered by environmental signals, MAPK kinase kinase, named as MAPKKK or MEKK, is firstly activated, then result in the phosphorylation of MAPK kinase, known as MAPKK, MEK or MKK, and finally MAPK was activated. In the end of the phosphorylation-activating steps, MAPKs bind with and phosphorylate amount of protein substrates to transferring upstream signals (Opdenakker et al., 2012, Meng and Zhang, 2013, Pitzschke, 2015, Xu and Zhang, 2015). As one well-studied MAPK, *Arabidopsis* MPK3 plays important roles in numerous processes, including defense against pathogens (Andreasson and Ellis, 2010, Galletti et al., 2011, Meng et al., 2012, Pitzschke, 2015). Tobacco MPK3 (NtMPK3), also known as SIPK, is an ortholog of *Arabidopsis* MPK3, which is reported to be activated in response to fungal infection and increase production camalexin (Ren et al., 2008).

The closest protein homologue of RSG in *Arabidopsis thaliana* is VIP1, which is known to interact with *Agrobacterium* effector VirE2 and mediated nuclei transport of T-complex during *Agrobacterium* mediated genetic transformation (Tzfira et al., 2001). Interestingly, VIP1 was shown to interact with *Agrobacterium*-induced mitogen-activated protein kinase (MAPK) MPK3, and VIP1 relocalizes from cytoplasm to nucleus upon phosphorylation by MPK3. Djamei et al also reported that MAPK-dependent phosphorylation of VIP1 is necessary for *Agrobacterium* mediated T-DNA transformation, and flg22, as a microbe associated molecular pattern, is able to cause VIP1 to relocaolized to the nuleus (Djamei et al., 2007). Moreover, a DNA hexamer motif named VIP1 response elements (VREs) was identified. By binding to VREs, VIP1 regulates the expression of MPK3 pathway related genes, such as *Trxh8* and *MYB44* (Pitzschke et al., 2009). Additionally, as a bZIP transcriptional factor in *thaliana,* VIP1 and it close homologues also showed liable localization in plant cells under abiotic stresses, such as hypo-osmotic stress, which mimic touch stimuli (Tsugama et al., 2016, Tsugama et al., 2014).

Considering the complexity of crosstalk throughout plant hormone pathways, it is meaningful to see if RSG also interacts with other protein kinases which are involved in different pathways, for example, MPK3 in tobacco. In this study, we identified a tobacco MPK3 (NtMPK3) as a RSG interaction protein kinase. By mutating potential MAPK associated phosphorylation sites in RSG, NtMPK3-RSG interaction complex displayed different intracellular localization. Triple mutation in all three MAPK associated phosphorylation sites (Thr-28, Ser-74 and Thr-135) to alanine completely blocked nuclear localization of RSG in tobacco cells. We also report on the role of MAPK associated phosphorylation sites as key regulators of RSG localization and its response to biotic or abiotic stresses. This work gives insight into the molecular mechanism whether MAPK pathway is linked to other plant hormone pathway regulators.

## Results

### *Bacillus cereus* AR156 induces nuclear transport of RSG

*B.cereus* AR156 is an effective biocontrol agent, which was previously isolated from natural environment (Jiang et al., 2017). *B. cereus* AR156 facilitates plant defense against variety of diseases through SA and JA/ET dependent pathways, which was known as Induced Systematic Resistance (ISR) (Niu et al., 2011). It was also reported that *B. cereus* AR156 treatment could induce the MPK3/MPK6 pathway in *Arabidopsis* by biosynthesizing extracellular polysaccharides (EPS) (Jiang et al., 2016a). The initial aim of this study was to investigate if biocontrol agents, such as *B. cereus* AR156, also manipulate nuclear localization of functional transcriptional factors in either plant defense or growth process. In previous research presented by Armin *et al., Arabidopsis* VIP1 relocalizes from cytoplasm to nucleus and regulates the expression of defense-related genes, such as *PR1*, at the present of flg22 or *Agrobacterium* strain (Djamei et al., 2007). It was further revealed that VIP1 binds with specific DNA segments and promote the expression of *Trxh8 and MYB44* to induce MPK3 pathway (Pitzschke et al., 2009). As an ortholog of VIP1 in tobacco, RSG shows 48% exact (177/370) and 56% positive (210/370) hits with *Arabidopsis* VIP1 in protein sequence alignment, and both RSG and VIP1 have a conserved basic leucine zipper (bZIP) domain (Figure. 1). By directly applying *B. cereus* AR156 suspension on transgenic tobacco which transiently expresses RSG with CFP tag, RSG started to relocalized in tobacco nuclear in 5 min, and pre-dominant nuclear localization was observed in 30 min (Figure. 2). This results indicated that not only the CDPK1 associated pathway, but also the MAPK mediated phosphorylation might be potentially involved in intracellular relocalization of RSG.

**Figure 1.**
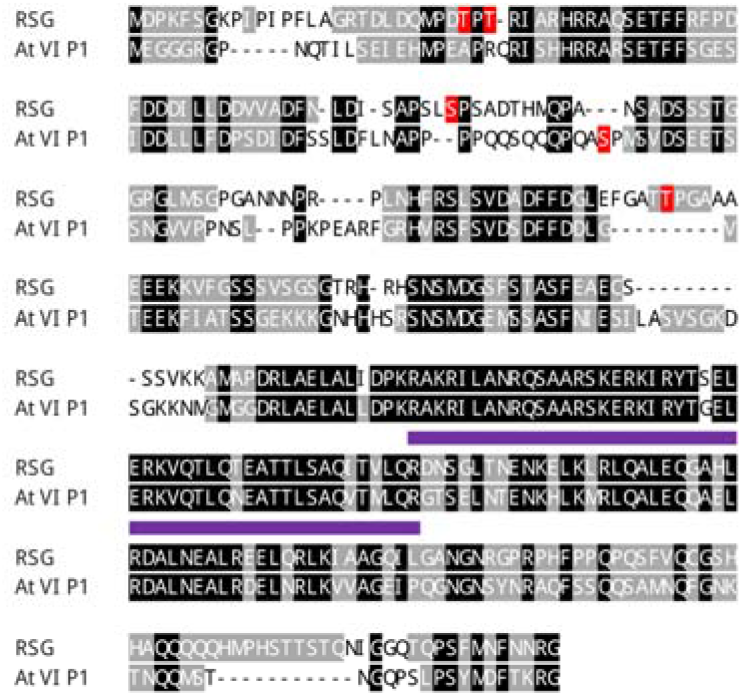
Amino acid sequence alignment of RSG and AtVIP1. The RSG from N. tabacum cv. Turk (accession number BAA97100.1) was aligned using the T-Coffee program (http://www.ebi.ac.uk/Tools/t-coffee/index.html) with its closest homolog AtVIP1 from Arabidopsis thaliana (AtVIP1, accession number NP_564486.1). Amino acid residues identical between RSG and AtVIP1 are highlighted in black and gray for conserved hits. The basic leucine zipper (bZIP) domains are underlined in purple. Amino acids potentially phosphorylated by MAPK are highlighted in red.

**Figure 2.**
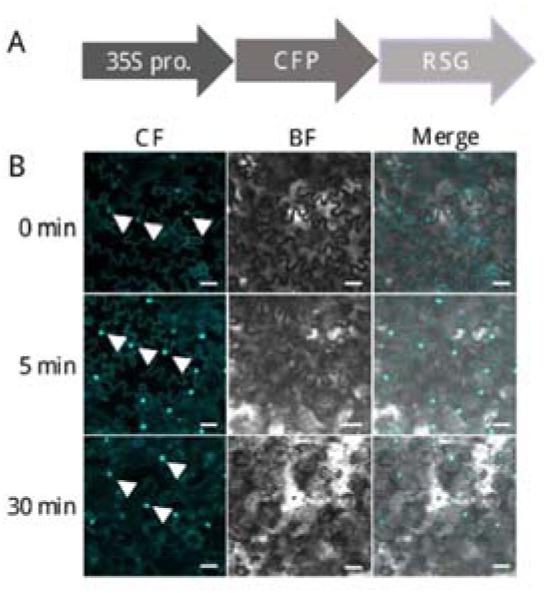
*B. cereus* AR156 induces nuclear import of RSG in *N. tabacum* cells. (A) Constructs of 35S-Figure 2. B. cereus AR156 induces nuclear import of RSG in N. tabacum cells. (A) Constructs of 35S-driven RSG-CFP. (B) Confocal microscopy observation of transgenic tobacco overexpresses RSG-CFP upon treatment of B. cereus AR156 suspension in 5 min and 30 min. RSG tagged with CFP were stably expressed in transgenic N. tabacum, and analyzed by confocal microscopy after B. cereus AR156 treatment. CFP signal is in cyan. Images are single confocal sections, representative of images obtained in three independent experiments. Scale bars= 40 μM. White arrows indicate observed nuclear of tobacco cells. Scale bars = 20 μm. Three independent experiments were performed for each assay with similar results.

### NtMPK3 interacts with RSG as a potential RSG kinase

MAPKs, as terminal protein kinases in plant mitogen-activated protein kinase cascades, directly interact with targeted substrate proteins and continue with phosphorylation process (Xu and Zhang, 2015). To verify the relationship between RSG and MAPKs in planta, we first investigate the interaction between RSG and NtMPK3 (XP_016478177.1), which was considered as one of representative tobacco MAPKs in this study. As shown in Figure 3A and 3B, we co-expressed RSG fused with nCerulean and NtMPK3 fused with cCFP in *N. benthamiana* leaves under the control of cauliflower mosaic virus 35S promoter (CaMV 35S), and interaction between RSG and NtMPK3 was observed in 3 days after *A. tumefaciens-*mediated infiltration. By using DsRed fused with previously identified nuclear localization site (NLS) of VirD2 (Jayaswal et al., 1987) as a nuclear indicator, RSG-NtMPK3 interaction complex showed a nuclear localizing pattern in *N. bentamiana* cells (Fig. 3B).

**Figure 3.**
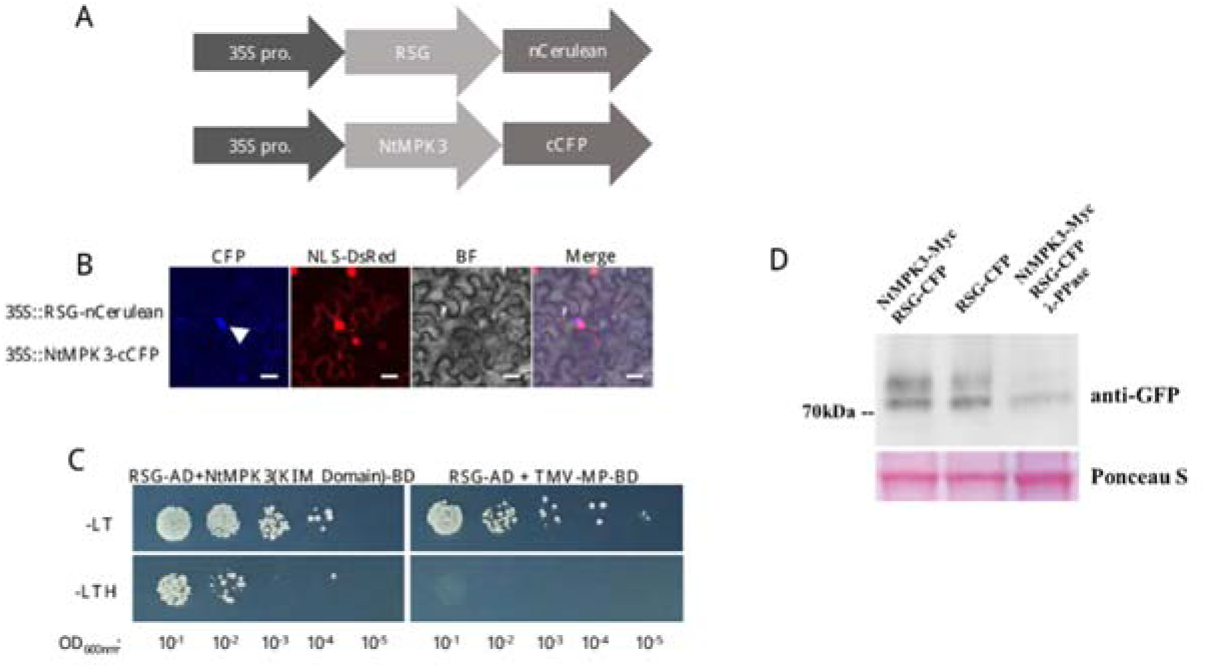
RSG interacts with mitogen-Activated Protein Kinase 3 (MAPK3) in tobacco. (A) Constructs of 35S-driven RSG-nCerulean and 35S-driven NtMPK3-cCFP. (B) BiFC assay. RSG-nCerulean and NtMPK3-cCFP were transiently expressed in agroinfiltrated leaf epidermis of N. benthamiana. NLS-VirD2 fused with DsRed indicated the localization of nuclear. and analyzed by confocal microscopy three days post-infiltration. CFP signal is in cyan. Images are single confocal sections, representative of images obtained in two independent experiments performed for each protein; for each experiment, three infiltrations were performed on three different leaves, with two images recorded per infiltration. Scale bars= 20 μM. Yeast-two-hybrid interaction assay. LexA-NtMPK3(KIM Domain) was coexpressed with Gal4-AD fused to RSG. The indicated dilutions of cell cultures were plated and grown on non-selective (+ histidine) and selective media (- histidine). Detection of phosphorylation of RSG coexpresses with NtMPK3. For λ-PPase treatment, 10 units μL-1 λ-PPase was added to the reaction and incubated at 30 °C for 30 min.Three independent experiments were performed for each assay with similar results.

Yeast-two-hybrid assay was then conducted to confirm the interaction between NtMPK3 and RSG in yeast cell system. As NtMPK3 showed irrepressible self-activation manner in yeast-two-hybrid assay, the kinase interaction motif (KIM) docking site domain (91-342aa) of NtMPK3 was identified and cloned into pSTT91. We expressed KIM domain of NtMPK3 fused with LexA-BD and RSG fused with GAL4-AD in *Saccharomyces cerevisiae* strain TAT7 (L40-ura3), in which cell growth on a histidine-deficient medium indicates interaction. Fig. 3C shows that KIM docking site domain of NtMPK3 interact with RSG, and interaction ability of RSG was specific because it was not observed with the cell-to-cell movement protein (MP) of the *Tobacco mosaic virus* (TMV), which was considered as an unrelated control protein. It is also revealed that KIM docking site domain of NtMPK3 was the only specific domain take charge of binding protein substrate, such as RSG (Fig. S1).

BiFC and yeast-two-hybrid results were then validated by an independent approach, in which GFP tagged NtMAPK3 was coexpressed in *N. benthamiana* leaves with c-Myc tagged RSG, isolated total protein was immunoprecipitated with anti-GFP antibody followed by western blot anaylsis. Fig. 4D shows that collected immunoprecipitates include both NtMPK3 detected by anti-GFP and RSG detected by anti-Myc, which confirms the interaction between RSG and NtMPK3 in yeast-two-hybrid system and BiFC assay.

Previous studies proved that AtVIP1 is a MAPKs associated protein substrate in *A. thaliana* (Djamei et al., 2007). To determine whether RSG, as an orthologue in tobacco, is a protein substrate of NtMPK3, we isolated protein extract from *N. tobacum* plant overexpresses RSG-CFP fusion protein, and western blot analysis with monoclonal anti-GFP antibody detected additional shifted bands (Fig. 3D). λ-protein phosphatase (λ-PPase) treatment decreased the relative intensity of shifted bands, which indicates that these shifted bands correspond to the phosphorylated status of RSG-CFP fusion. Collectively, these results in Figure 3 support the notion that RSG not only interacts with NtMPK3 both in vitro and in vivo, but also is a protein substrate of NtMPK3 during phosphorylation process.

### MAPK associated phosphorylation sites in RSG regulates its intracellular localization with NtMPK3

Previous study identified Ser-114 in RSG, which was a phosphorylation site of a NtCDPK1, manipulates the interaction between 14-3-3 and RSG (Ishida et al., 2008). Their results suggested that impaired phosphorylation on Ser-114 of RSG results in the reduction of expression of RSG-targeted genes (Igarashi et al., 2001, Ishida et al., 2004). To determine whether MAPK associated phosphorylation sites of RSG were also involve in its downstream biological functions, we generated a set of potential mutant versions of RSG in which MAPK associated phosphorylation sites was altered from Ser or Thr to Ala to simulating durable non-phosphorylation status, which were recorded as RSG^T28A^, RSG^T30A^, RSG^S74A^ and RSG^T135A^ (Fig. 4) Interestingly, all of these generated RSG mutations showed interaction with KIM docking site domain of NtMPK3 in yeast-two-hybrid assay, while only RSG^T28A^ still co-localized with NtMPK3 in nucleus of *N. benthamiana* cell (Fig. 4A and 4B).

**Figure 4.**
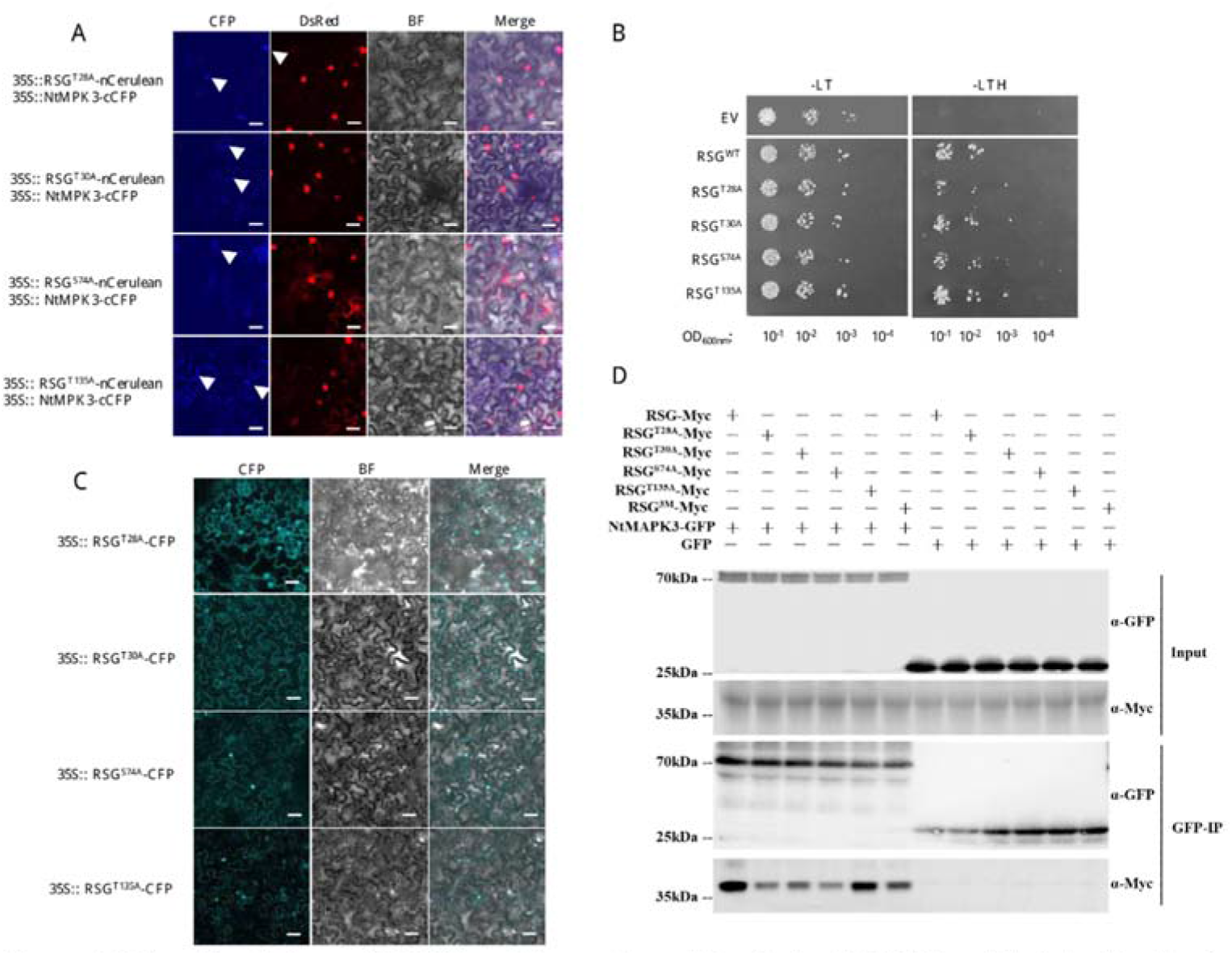
Interaction between NtMPK3 and four amino acid mutants of NtRSG and their localization in tobacco cells by BiFC. (A) BiFC assay. RSG-nCerulean or RSG derived single point amino acid mutants fused with nCerulean and NtMPK3-cCFP were coexpressed in N. benthamiana. NLS-VirD2 fused with DsRed refer to the localization of nuclear. Scale bars = 20 μm. (B) Yeast-two-hybrid interaction assay. LexA-NtMPK3(KIM Domain) was coexpressed with Gal4-AD fused to RSG or RSG. The indicated dilutions of cell cultures were plated and grown on non-selective (+ histidine) and selective media (- histidine). (C) RSG and indicated mutants tagged with CFP were transiently expressed in agroinfiltrated leaf epidermis of N. benthamiana, and analyzed by confocal microscopy three days post-infiltration. CFP signal is in cyan. Images are single confocal sections, representative of images obtained in two independent experiments performed for each protein; for each experiment, three infiltrations were performed on three different leaves, with two images recorded per infiltration. Scale bars= 40 μM. (D) Co-immunoprecipitation interaction assay. NtMAPK3-GFP was expressed with C-terminus Myc tagged RSG and its relative mutants for 3 days in agroinfiltrated Nicotiana benthamiana leaves, immunoprecipitated (IP) with anti-GFP antibody (top panel), followed by western blot analysis with anti-GFP or anti-Myc antibody. Two independent experiments were performed for each assay with similar results.

Unlike their co-localization with NtMPK3, RSG mutants displayed both nuclear and cytoplasm localization in *N. benthamiana* cells by agro-infiltration (Fig. 4C). However, nuclear localizations of RSG^T30A^-CFP, RSG^S74A^-CFP and RSG^T135A^-CFP were lackluster, although RSG^T28A^-CFP showed brighter nuclear localization (Fig. 4C). Consequently, Thr-30, Ser-74 and Thr-135 might play more important role during NtMPK3 associated phosphorylation of RSG.

### RSG^T30A^ and RSG^S74A^ show durable localization in tobacco nuclear after *B. cereus* AR156 treatment

*Agrobacterium* treatment was reported to induce nuclear localization of AtVIP1 in *Arabidopsis* cells (Djamei et al., 2007). Thus, transgenic tobacco lines which stably express RSG^T30A^-CFP and RSG^S74A^-CFP driven by 35S promoter were generated to avoid possible biotic stimulation from *Agrobacterium* during agro-infiltration. Differ from observed results in Fig. 4C, stably expressed RSG^T30A^-CFP and RSG^S74A^-CFP displayed mere cytoplasm localization in *N. tobacum* cells, in which *Agrobacterium* stimulation was excluded (Fig. 5).

**Figure 5.**
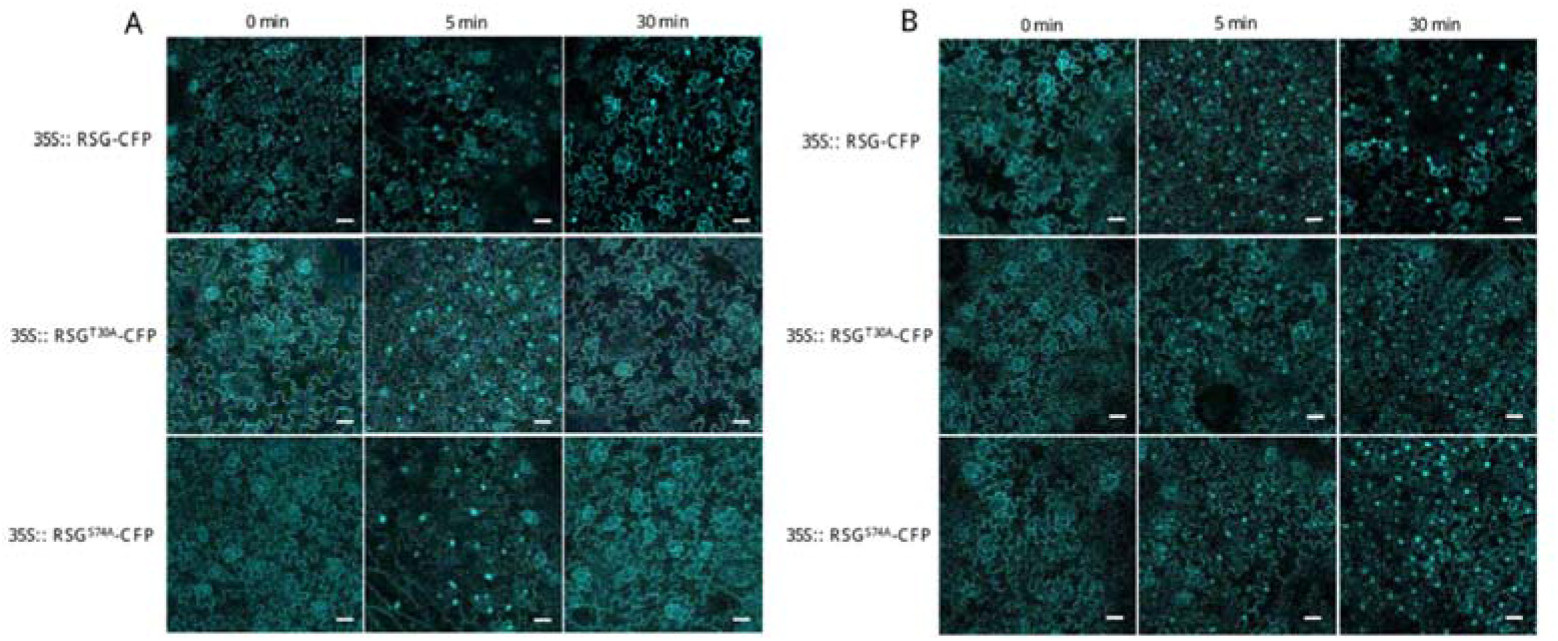
RSG^T30A^ and RSG^S74A^ show durable localization in tobacco nuclear after *B. cereus* AR156 treatment. (A) Observation of transgenic tobacco overexpressed RSG, RSG^T30A^ or RSG^S74A^ fused with CFP upon treatment of flg22. RSG and indicated mutants tagged with CFP were stablely expressed in transgenic *N. tabacum*, and analyzed by confocal microscopy after flg22. (B) Observation of transgenic tobacco overexpressed RSG, RSG^T30A^ or RSG^S74A^ fused with CFP upon treatment of *B. cereus* AR156 suspension. RSG and indicated mutants tagged with CFP were stablely expressed in transgenic *N. tabacum*, and analyzed by confocal microscopy after *B. cereus* AR156 treatment. CFP signal is in cyan. Images are single confocal sections, representative of images obtained in two independent experiments performed for each protein; for each experiment, three infiltrations were performed on three different leaves, with two images recorded per infiltration. Scale bars= 40 μM.

To investigate whether NtMPK3 activation could affect the intracellular localization of RSG in tobacco cells, flg22, a well-studied peptide corresponding to conserved flagellin sequence of pathogenic bacteria that initiate plant MAPK signaling cascade (Djamei et al., 2007), and *B. cereus* AR156 cell suspension were applied on acquired transgenic tobacco leaves (Fig. 5A and B). Upon treatment of flg22, we observed strong increased signal of RSG-CFP, RSG^T30A^-CFP and RSG^S74A^-CFP in the tobacco nuclear in 5 min. However, RSG^T30A^-CFP and RSG^S74A^-CFP relocalized in cytoplasm in 30 min, whereas RSG-CFP remained significant nuclear localization (Fig. 5A). Interestingly, *B. cereus* AR156, as an effective plant defense elicitor, facilitated lasting nuclear transport of RSG-CFP, RSG^T30A^-CFP and RSG^S74A^-CFP in 30 min, which might due to that *B. cereus* AR156 provides more intensive and continuous stimulations to plant than flg22 (Fig. 5B). These results support the notion that MAPK associated phosphorylation sites of RSG are also involved in its intracellular relocalization.

### Mimic constant MAPK associated unphosphorylation in RSG leads pre-dominant cytoplasm localization

As shown in previous research in VIP1, Ser-79, a MPK3 targeted phosphorylation site was mutated to Asp to obtain constant phosphorylated VIP1, and this VIP1 mutation showed pre-dominantly (Djamei et al., 2007). In contrast to preventing phosphorylation on MAPK associated sites in RSG, Thr-30, Ser-74 and Thr-135 in RSG were mutated to alanine to mimic constitutively nonphosphorylated status, which was marked as RSG^3M^. Interestingly, RSG^3M^ shows pre-dominantly cytoplasm localization in *N. benthamiana* by agro-infiltration (Fig. 6A). To better evaluate the difference between RSG^3M^ and other RSG mutants and exclude the stimulation from *Agrobacterium*, we generate transgenic *N. tobacum* overexpressed RSG^3M^ fused with CFP tag. As shown in Fig. 6B, RSG^3M^ naturally localized in tobacco cytoplasm, and no significant nuclear relocalization was observed although flg22 and *B. cereus* AR156 were applied.

**Figure 6.**
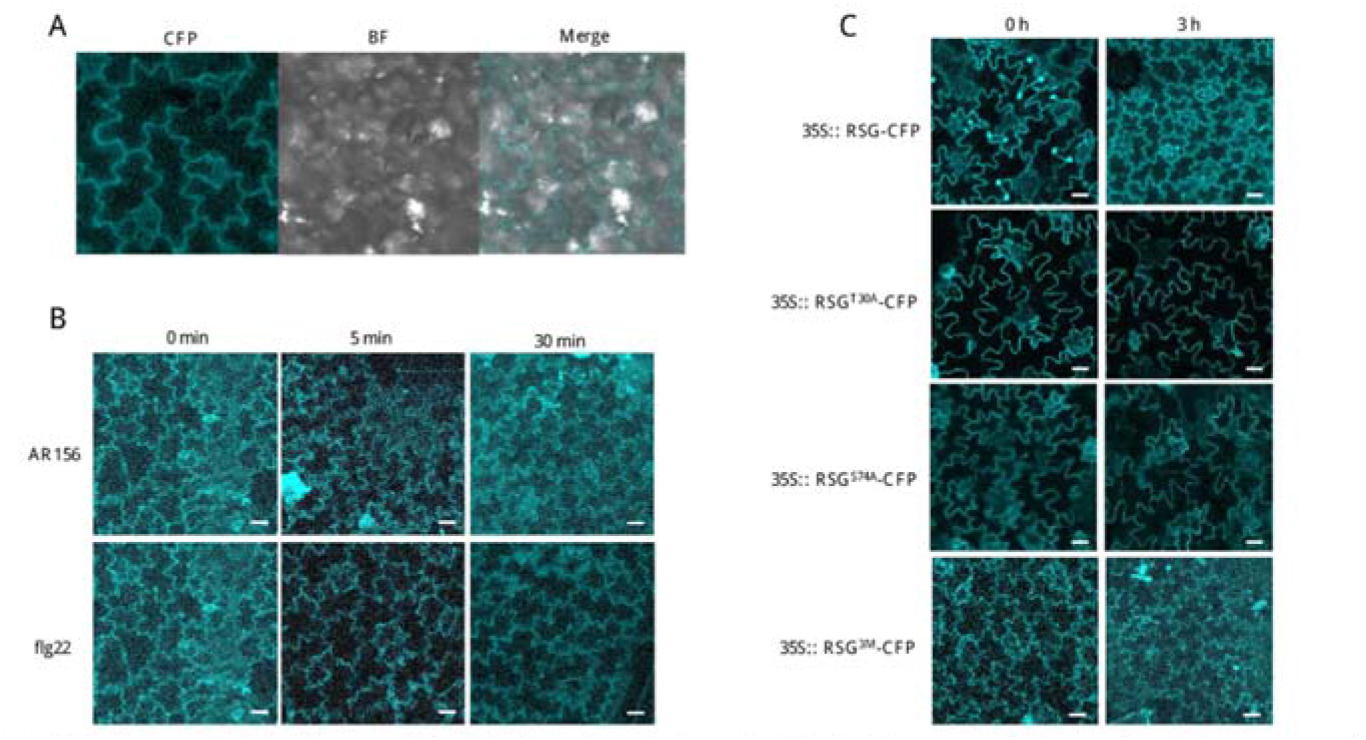
Mimic non-MAPK associated phosphorylation in RSG shows stable pre-dominant cytoplasm localization. (A) RSG 3M (T30D/S74D/T135D) tagged with CFP were transiently expressed in agroinfiltrated leaf epidermis of *N. benthamiana*, and analyzed by confocal microscopy three days post-infiltration. (B) RSG^3M^ (T30A/S74A/T135A) tagged with CFP were stably expressed in transgenic *N. tabacum*, and analyzed by confocal microscopy upon treatment of flg22 or *B. cereus* AR156 suspension. (C) Observation of transgenic tobacco overexpressed RSG, RSG^T30A^,RSG^S74A^ or RSG^3M^ fused with CFP upon treatment GA3. RSG and indicated mutants tagged with CFP were stablely expressed in transgenic *N. tabacum*, and analyzed by confocal microscopy 3 hours after GA_3_ treatment. CFP signal is in cyan. Images are single confocal sections, representative of images obtained in two independent experiments performed for each protein; for each experiment, three infiltrations were performed on three different leaves, with two images recorded per infiltration. Scale bars= 40 μM.

Ishia et al reported that RSG transports to plant nucleus in response to the reduction of plant GA contents, and exogenous GA treatment could lead the RSG cytoplasm relocalization (Ishida et al., 2004). Related study also revealed that CDPK1 dependent pathway is involved in the GA-induced RSG nuclear export, and phosphorylation of Ser-114 was specifically important in this process (Ishida et al., 2008). In our study, apparent nuclear export was also observed 3 hours after GA3 treatment in transgenic tobacco which overexpresses RSG-CFP (Fig. 6C). RSG mutants, including alanine substitution of Thr-30 or Ser-74 and alanine substitution of Thr-30, Ser-74 and Thr-135, showed stable pre-dominant cytoplasm localization both in pre- and post-GA treatment (Fig. 6C). Collectively, this part of results has identified a triple amino acid mutant of RSG which maintains stable cytoplasm intracellular localization, and MAPK associated phosphorylation sites in RSG might also participate in GA regulated RSG nuclear export in tobacco cells.

Constant MAPK associated phosphorylation in RSG reduces expression of downstream genes.

RSG is also a functional transcriptional factor in tobacco, and a GA biosynthetic pathway related gene *NtGA20ox1* was reported to be a direct target of RSG (Fukazawa et al., 2010). To examine whether MAPK associated phosphorylation sites of RSG regulate its function as transcriptional factor in GA biosynthetic pathway, we analyzed the expression level of *NtGA20ox1* in transgenic tobacco plants which overexpress RSG or RSG mutants (Fig. 7A and B). By Semi-quantitative PCR assay, the highest expression level of *NtGA20ox1* was detected in RSG overexpressed tobacco, and the lowest expression of *NtGA20ox1* was detected in RSG^T30A^, RSG^S74A^ and RSG^3M^ overexpressed tobacco (Fig. 7A and B). These results in Fig. 7A and B are consistent with presented data in Fig. 5A and Fig. 6A, in which alternations of MAPK associated phosphorylation sites in RSG significantly lead to its pre-dominant cytoplasm localization.

**Figure 7.**
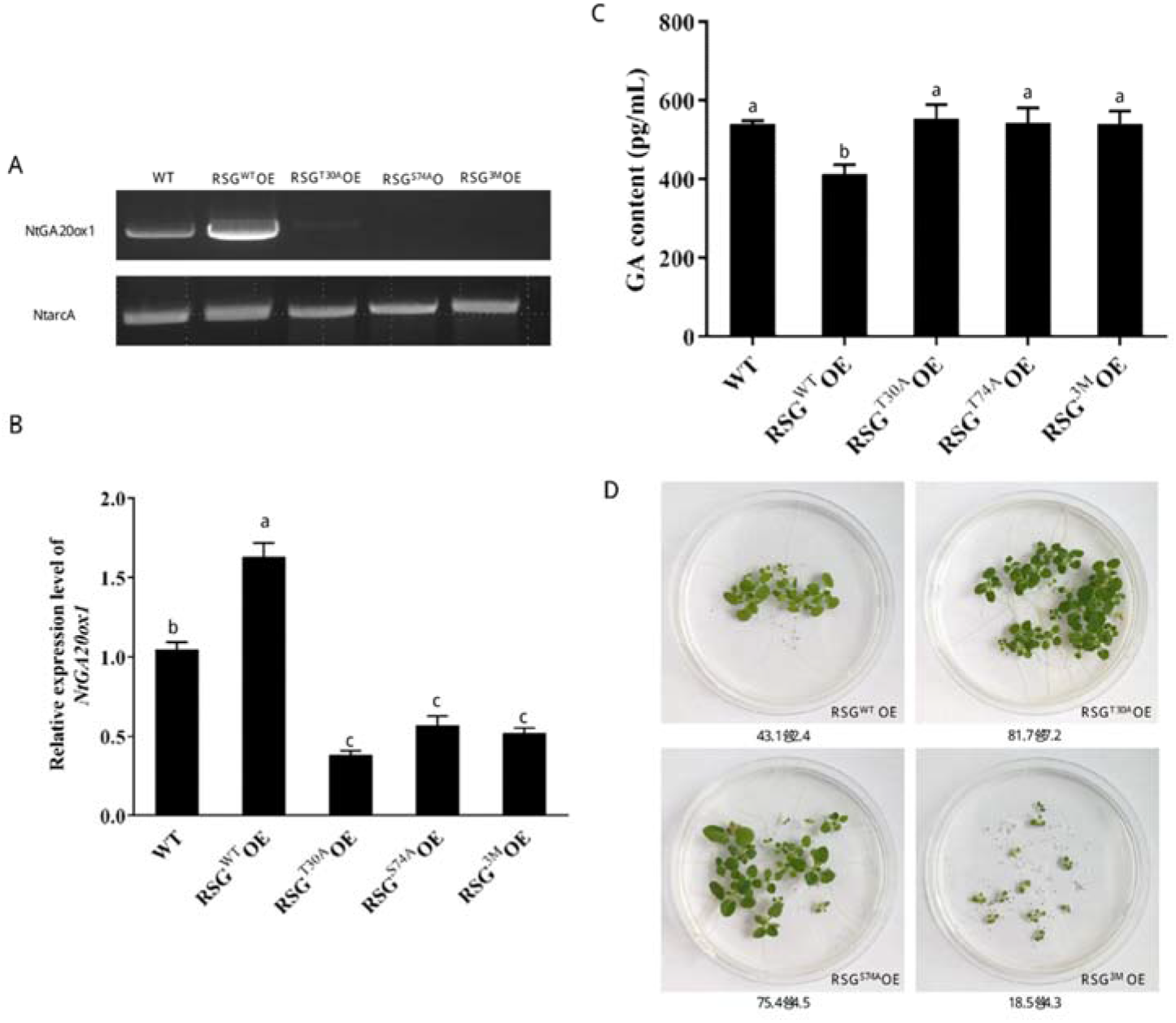
MAPK associated phosphorylation sites in RSG regulates its function as transcriptional factor. (A)-(B) Semi-quantitative PCR analysis of *NtGA20ox1* expression in transgenic tobacco. (C) Detection of GA contents in transgenic tobacco overexpresses RSG or its related mutants. The values are Mean ± Standard Deviation, followed by the same letter within a column are not significantly different as determined by the Duncan’s multiple range tests test (*P < 0.05*). (D) Seeds of transgenic tobacco overexpressing RSG or its relative mutants were germinated in MS medium. The values at the bottom of panels indicate the germination ratio calculated from three independent plates. Three biological replicates were performed for each assay with similar results.

We further compared the GA contents in acquired transgenic tobacco overexpressed RSG or its mutants with wild-type tobacco (Fig. 7C). Interestingly, RSG overexpressed tobacco maintains the lowest GA contents compared with wild-type tobacco and other transgenic tobacco plants, although RSG overexpressed tobacco showed the highest expression level of *NtGA20ox1* (Fig. 7C). This might due to large amount of RSG accumulation in tobacco nuclear could trigger negative feedback during GA biosynthesis. These results confirmed that MAPK associated phosphorylation sites in RSG not only regulate its intracellular localization, but also alter its biological functions, for example, as a transcriptional factor. Interestingly, geminating rate of transgenic tobacco seeds of as low as 43.1%, and recovered to 81.7% and 75.4% when Thr-30 or Ser-74 was mutated to alanine. The transgenic tobacco seeds overexpress RSG^3M^ only produces weak and growth aborting seedlings by 18.5% of germination rate (Figure 7D).

## Discussion

Tobacco transcription factor RSG was first identified in tobacco with function of regulating morphology of plants by controlling the endogenous amounts of Gas (Fukazawa et al., 2000). It was then discovered to control the feedback regulation of NtGA20ox1 via intracellular localization and epigenetic mechanism, and signaling protein 14-3-3 was proven to be RSG binding partner and participate in nuclear-cytoplasmic shuttling (Igarashi et al., 2001, Fukazawa et al., 2011). Tobacco Ca^2+^-dependent protein kinase CDPK1 (NtCDPK1) was identified as a RSG kinase which promotes 14-3-3 binding of RSG by phosphoraylation of RSG (Ishida et al., 2008, Ito et al., 2014). Our proposed involvement of the RSG interacting with MPK3 is consistent with VIP1, an *Arabidopsis* homologue of RSG, in previous research (Djamei et al., 2007). Interestingly, VIP1 showed several similarities with RSG: VIP1 presents nuclear-cytoplasmic shuttling subcellular localization pattern (Tsugama et al., 2012) and VIP1 interacts with 14-3-3 and its subcellular localization is regulated by phosphorylation (Takeo and Ito, 2017).

*Agrobacterium* is known to priming plant defense response (Sheikh et al., 2014), thus, most of the experiments in our study were carried on transgenic plants. In Figure 4C, subcellular localizations of RSG mutant were observed by *Agrobacterium* infiltration on *N. benthamiana* leaves, and both cytoplasm and nuclear localization was observed for RSG^T28A^, RSG^T30A^, RSG^S74A^ and RSG^T135A^. In transgenic tobacco, RSG^T30A^ and RSG^S74A^ shows only cytoplasm localization before flg22 and *B. cereus* AR156 treatment, indicates *Agobacterium* infiltration process actually stimulate nuclear transport of RSG protein and related mutants (Figure 5). HXRXXS motifs are demonstrated as 14-3-3 binding targets (Yaffe et al., 1997), and S114 in RSG, S35, S115, S151 in VIP1 were identified as HXRXXS motifs. The mutated RSG on S114 site to alanine cannot bind to 14-3-3 and exclusively localized in the nucleus, and the nonphosphorylated VIP1 mutant of three HXRXXS motifs showed no interaction with 14-3-3 and only localizes in nucleus (Igarashi et al., 2001, Takeo and Ito, 2017). Importantly, the four predicted MAPK mediated phosphorylation sites of RSG are not correspond to the pattern of HXRXXS motif (Figure 1), indicating blocking interaction with 14-3-3 might not be the cause of changes in nuclear-cytoplasmic localization of mutated RSG in this study. Moreover, artificial mutating MAPK associated phosphorylation sites of RSG to alanine results in only cytoplasmic protein localization, which was contrary to dominant nuclear localization of RSG mutated on NtCDPK1 associated site (Igarashi et al., 2001). One possible explanation is RSG, as an important bZIP transcription factor, might be phosphorylated by different protein kinase on different motifs under different stimulations, for example, abiotic and biotic stresses.

In our result, biotic stimulations, for example, bacterial flagellin peptide flg22, induces nuclear import of RSG up to 30 min (Figure 5A). However, on transgenic tobacco overexpresses RSG^T30A^ and RSG^S74A^, mutated RSG rapidly relocalized back to cytoplasm in 30 min post flg22 treatment, although nuclear import was observed in initial 5 min. Single mutation of MAPKs associated sites partly impaired nuclear import of RSG under biotic stimulation, as flg22 was proven to trigger activation of MAPK cascade in plant (Pitzschke et al., 2009). Biocontrol agent strain *B. cereus* AR156 is also reported to prime plant defense, and MAPK signaling pathway is involved in this process as well (Niu et al., 2011, Nie et al., 2017). Interestingly, continuous nuclear transport of RSG^T30A^ and RSG^S74A^ was detected on transgenic tobacco leaves treated with *B. cereus* AR156 in 30 min (Figure 5B). The reason might be, unlike flg22, *B. cereus* AR156 cells might provide efficient and constant biotic stimulations on treated plant tissues. Although one MAPK associated phosphorylation site was mutated, phosphorylation on the rest sites (Ser-74, Thr-135 in RSG^T30A^, and Thr-30, Thr-135 in RSG^S74A^) would still contribute to nuclear transport of RSG under biotic stresses, and stronger stimulation might trigger longer retention time of RSG in plant nuclear. Indeed, *B. cereus* AR156 was proven to prime plant defense by several ways, including secreting extracellular polysaccharides, manipulating specific miRNA or transcription factors in treated plant (Nie et al., 2017, Jiang et al., 2016a, Jiang et al., 2016b).

To better illustrate the importance of MAPK associated phosphorylation sites in RSG during nuclear-cytoplasmic shuttling under biotic stresses. We generated triple points mutant of RSG on three MAPK associated phosphorylation sites to alanine, including Thr-30, Ser-74 and Thr-135, and results in RSG^3M^. Consistent with single mutants RSG^T30A^ and RSG^S74A^, RSG^3M^ shows only cytoplasm localization in transgenic plant before treatment, while neither flg22 nor *B. cereus* AR156 treatment could induce nuclear transport of RSG^3M^ (Figure 7A and B). These data further manifest importance of MAPK associated phosphorylation sites of RSG in response to biotic stresses in plant.

The nuclear localization of RSG is directly related to its function as transcription factor. The downstream gene regulating capacity of RSG plays important role in GA feedback regulation in tobacco, and *NtGA20ox1* was reported as downstream gene regulated by RSG (Ishida et al., 2008, Fukazawa et al., 2011). Reduction of *NtGA20ox1* expression level in transgenic tobacco of RSG^T30A^, RSG^S74A^ and RSG^3M^ was consistent with predominant cytoplasm localization of these RSG mutants. As previous study demonstrated RSG could interact with itself like other plant bZIP family proteins, overexpression of point mutated version of RSG might hijack endogenous RSG and blocking its native functions (Fukazawa et al., 2000). However, we did not detect significant higher GA content in transgenic tobacco of RSG^T30A^, RSG^S74A^ and RSG^3M^ compared with wild type plant, which might because of the complicated plant hormone feedback regulating mechanisms. Specifically, germination of seeds obtained from RSG^3M^ transgenic tobacco was relative slow and incomplete compared with seeds obtained from RSG, RSG^T30A^ and RSG^S74A^ overexpression tobacco, suggests function of RSG as transcription factor is much more important for tobacco seed germinating stage.

Collectively, the interaction between RSG and NtMPK3, the involvement of MAPK associated phosphorylation sites in nuclear-cytoplasmic shuttling of RSG and observed phenotype variations in transgenic plants overexpress mutated version of RSG suggest that phosphorylation on MAPK associated sites on RSG deeply regulate its function in plant. Interestingly, RSG was previously proven to bind VIP1 response element (VRE) sequence and activate downstream gene expression (Wang et al., 2018), indicates potential target gene range overlaps between RSG and VIP1. It would be particularly interesting to investigate further functions of RSG in downstream defense response against plant pathogens.

## Conclusions

In our research, we found tobacco RSG is involved in gibberellin feedback regulation by inducing the expression of key genes and we identified tobacco mitogen-activated protein kinase 3 (NtMPK3) as a RSG interacted protein kinase. Mutation of predicted MAPK-associated phosphorylation site of RSG (Thr-30, Ser-74 and Thr-135) significantly altered the intracellular localization of NtMPK3-RSG interaction complex. Nuclear transport of RSG and its amino acids mutants (T30A and S74A) were observed after treated with plant defense elicitor peptide flg22 in 5 min, while the two mutated RSG swiftly relocalized in tobacco cytoplasm in 30 min. Moreover, triple points mutation of RSG (T30A/S74A/T135A) mimics constant unphosphorylated status, and predominantly localized in tobacco cytoplasm. RSG (T30A/S74A/T135A) showed no relocalization effect under the treatments of either flg22, *B. cereus* AR156 or GA_3_, and was impaired in its role as TFs. Our results suggest that MAPK associated phosphorylation sites of RSG regulate its localization in tobacco and constant unphosphorylation of RSG in Thr-30, Ser-74 and Thr-135 keeps RSG predominantly localized in cytoplasm.

## Material and methods

### Bacterial strains, plant and growth conditions

*A. tumefaciens* strains and *E. coli* strains were grown in on Luria–Bertani (LB) agar (NaCl 10g·L-1, Yeast extract 5g·L-1, Tryptone 5g·L-1) at 28 °C. *B. cereus* AR156 was isolated from the forest soil of Zhenjiang City, Jiangsu Province, China, as an effective bacterial BCA (Genebank accession number CP015589), grown in LB medium at 30 °C. *Nicotiana tabacum* var. Turk and *Nicotinana bentaminan* plants were grown in soil or in MS medium (MES 0.5 g/L, sucrose 30 g/L, agar 8 g/L, pH5.8) after seed surface sterilization, and maintained in vitro. All plants were grown in environment-controlled growth chambers under long-day conditions (16 h light/8 h dark cycle at 140 μE sec^-1^m^-2^ light intensity) at 22 °C.

### Plasmid construction and mutagenesis

Primer sequences used in these cloning procedures are described in Table S2, and plasmids and cloning strategies are summarized in Table S1. For Gal4-AD fusions, the coding sequences of RSG or RSG mutant variants were PCR-amplified, using total *N. tabacum* cDNA library as template, and cloned into the indicated sites of pGAD424 (LEU2+, Clontech; Mountain View, CA). For LexA fusions, the coding sequences of NtMPK3 and AtMPK3 were PCR-amplified, using total *N. tabacum* cDNA library as template, and cloned into the indicated sites of pSTT91 (TRP1+) (Sutton et al., 2001). For generating point mutations in RSG, overlapping PCR reactions were conducted to generate codon substitutions. Two DNA segments were amplified by PCR: from the translation initiation codon at the 5’-end to the target codon position and from the target codon position to the 3’ end of the coding sequence. For example, to generate the T28A mutation in RSG, PCR reactions were performed with the primer pairs 8F-17R and 17F-8R (Table S). The two PCR products then were used as template to generate the full length mutated RSG sequence by overlapping PCR with the primer pair, followed by insertion of the resulting PCR product, that encodes the full-length RSG T28A mutant, into the BamHI/PstI site of plasmid pGAD424, resulting in pGAD424-RSG-T28A (Table S). Aiming to expressing targeted proteins in plants, the coding sequences of RSG was inserted into the multiple cloning sites of pSAT5-CFP-C1 for subcellular localization experiments or pSAT1-ncerulean-C1/pSAT5-cCFP-C1 for BiFC assays (Tzfira et al., 2005). Resulting pSAT series expression cassettes were excised with AscI for pSAT1 or I-CeuI for pSAT5 and transferred into the same site of the binary pPZP-RCS2, or pPZP-DsRedNLS-RCS vector with DsRed signal fused with a NLS tag of VirD2 (Ballas and Citovsky, 1997).

### Yeast-two-hybrid protein interaction assay

For yeast-two-hybrid experiments, the pSTT91 and pGAD424 plasmids cloned with potential interactors were introduced into the Saccharomyces cerevisiae strain L40 and grown for 2 days at 30°C on a leucine- and tryptophan- deficient synthetic defined premixed yeast growth medium (SD-Leu-Trp, TaKaRa Clontech)(Hollenberg et al., 1995). Five to ten acquired yeast colonies on SD-Leu-Trp plates were resuspended in sterilized water and plated on SD-Leu-Trp and same medium plate lacking leucine, tryptophan and histidine (SD-Leu-Trp-His). Yeast cell growth was observed after incubation at 30°C for 2-3 days.

### Biomolecular fluorescence complementation assay

For biomolecular fluorescence complementation (BiFC) assay, tested protein pair were fused with either nCerulean or cCFP tag and cloned into pSAT series plasmid (Tzfira et al., 2005). For example, to observe the interaction between RSG and NtMPK3, the coding sequence of RSG was fused with nCerulean in pSAT1-nCerulean-C1, the coding sequence of NtMPK3 was fused with cCFP in pSAT5-cCFP-C1. The resulting expression cassette of RSG was then inserted into the pPZP-RCS1 binary vector and the expression cassette of NtMPK3 was inserted into the pPZP-RCS1-DsRedNLS binary vector. These two constructs were transiently co-expressed in *N. benthamiana* by agroinfiltration. CFP and DsRed fluorescence signal were detected after 2 days with a CLSM (Leica AF6000 modular microsystems). All experiments were biologically repeated for three times.

### Semi-quantitative PCR

For semi-quantitative PCR assays, total RNA from the shoots of transgenic tobacco expressing RSG-CFP or its related mutations under the control of CaMV 35S promoter or control wild-type *N. tabacum* were converted into cDNA with PrimeScript^TM^ RT reagent Kit (TaKaRa). PCR was performed with cDNA derived from 0.5 μg of total RNA with Extaq (TaKaRa). The primer sequences were 5’-CAACGCCCATCGTTTCATGG-3’ and 5’-CAAAAACTTGAAGCCCGCCA-3’ for NtGAox20 and 5’-ATGTGTTCGTTTCAGCCCGA-3’ and 5’-CCGCTGAAAAGTGTGCTTCC-3’ for tobacco *arcA* as an internal gene control (Ishida et al., 2004).

### Gibberellin content assay

Endogenous gibberellin (GA) contents were detected by Plant GA ELISA Kit (FEIYA Biotechnology) with double antibody sandwich method. Specifically, microwell plate were first coated by purified GA antibody, plant tissue extraction and GA-HRP antibody were then added into the wells for 1 h antibody staining. After drastically washing, TMB (3,3′,5,5′-tetramethylbenzidine) was added into wells as reaction substrate. The color of reaction solution was transformed to yellow in the presence of HRP at low pH value, and the OD reads at 450nm represent GA contents by referring to the standard curve.

### Confocal fluorescence microscopy

CLSM (Leica AF6000 modular microsystems) was used to collect sample images. A 434-nm line from an argon ion laser were used to excite cyan fluorescent protein (CFP), a 558-nm line from an argon ion laser were used to excite Red fluorescent protein (DsRed). For each assay, six independent leaves were observed for each experiment and each experiment has at least repeats.

### Statistical analysis

One-way analysis of variance (ANOVA) was carried out with SPSS (Version 19.0) and followed with Duncan’s multiple range tests (P < 0.05) for statistically analyzed and Student’s t test for evaluating the significance in all data.

## Acknowledgement

This research was supported by the National Natural Science Foundation of China (31900303, 31701829, 31972322), the Science and Technology Project of Jiangsu Province (BE2020408, BE2021364), the China Agriculture Research System of MOF and MARA (CARS-21), the Opening Project of Key Construction Laboratory of Probiotics in Jiangsu Province (JSYSZJ2019003)

---

**Figure S1.** Yeast-two-hybrid assay between 3 potential substrate binding domains of NtMPK3 and NtRSG. A. pSTT91-NtMPK3(Acitve site 48-246aa), pGAD424-TMV-MP. B. pSTT91-NtMPK3 (Acitve site 48-246aa), pGAD424-RSG ; C. pSTT91-NtMPK3 (Polypeptide substrate binding site 90-251aa), pGAD424-TMV-MP. D. pSTT91-NtMPK3(Polypeptide substrate binding site 90-251aa), pGAD424-RSG. E. pSTT91-NtMPK3(KIM domain 91-342aa), pGAD424-TMV-MP. F. pSTT91-NtMPK3(KIM domain 91-342aa), pGAD424-RSG.

**Figure S2.** RSG interacts with Arabidopsis mitogen-Activated Protein Kinase 3 (MPK3). (A) Constructs of 35S-driven RSG-nCerulean and 35S-driven AtMPK3-cCFP. (B) BiFC assay. RSG-nCerulean and AtMPK3-cCFP were transiently expressed in agroinfiltrated leaf epidermis of *N. benthamiana.* NLS-VirD2 fused with DsRed indicated the localization of nuclear. and analyzed by confocal microscopy three days post-infiltration. CFP signal is in cyan. Images are single confocal sections, representative of images obtained in two independent experiments performed for each protein; for each experiment, three infiltrations were performed on three different leaves, with two images recorded per infiltration. Scale bars= 20 μM.

**Table S1.** Plasmids and cloning strategies.

**Table S2.** Primer sequences.

